# Culsma: An Executable Specification Language for Laboratory Protocols

**DOI:** 10.64898/2026.05.07.723509

**Authors:** Meng Sun, Tadepally Lakshmikanth, Jun Wang, Hugo Barcenilla, Laura González, Yang Chen, Petter Brodin

**Affiliations:** Department of Women’s and Children’s Health, Karolinska Institutet, Stockholm, Sweden; Department of Immunology and Inflammation, Imperial College London, London, United Kingdom

## Abstract

Laboratory protocols are commonly communicated in natural language, leaving room for variation in how experimental commitments are understood and carried out. We present Culsma, an executable specification language in which protocols are written as programs. The same program is legible to bench scientists and directly machine-readable by software that checks and runs it. It serves as the protocol’s operational specification, making experimental commitments explicit for checking before execution and retaining them in results. Culsma addresses how experimental procedures are modeled: it defines operations by the experimental changes and relationships they establish, giving a fine-grained, compositional basis whose technique- and device-specific realizations remain extensible. We examine this boundary in workflows spanning molecular and cell biology, protein biochemistry, and immunoassays, which combine material handling, separation, and measurement differently, asking which distinctions the operation interface preserves. A maintained, source-linked benchmark broadens this evaluation with versioned checks of source-step correspondence and software execution.

## 1 Introduction

Laboratory protocols remain difficult to execute consistently across teams, time, and instruments. In most wet-lab settings, a protocol is still written as prose, interpreted by a human operator, and then translated again into device-specific actions, spreadsheets, or local checklists. Although human-readable, prose often leaves experimental commitments implicit, context-dependent, or open to interpretation. Step order, intermediate material identity, separation outputs, readouts, and handling requirements may depend on local knowledge rather than the written procedure itself. Consequently, the same written protocol can be interpreted differently across researchers and laboratories, before any question of programmatic execution arises [1, 2].

This limitation becomes more consequential as artificial intelligence and laboratory automation converge on iterative modes of experimentation [3]. In areas including systems biology, synthetic biology, and AI-assisted drug discovery, computational systems increasingly analyze experimental data, propose follow-up conditions, and pass them to automated execution platforms; the resulting observations can then inform the next round of analysis. Such cycles can expand experimental throughput and shorten the path from observation to follow-up experiment. Their operation, however, depends on a protocol that preserves experimental intent as it moves between scientists and software. A bench scientist must be able to inspect and modify the procedure, while computational systems need the same procedure in an operational form that they can validate and execute. Without such a shared representation, repeated protocol translation remains a bottleneck for AI-integrated experimentation.

This paper presents *Culsma*, an executable specification language in which laboratory protocols are written as programs. A Culsma source program serves as the protocol’s operational specification, expressing its experimental commitments in a form that can be read by scientists and interpreted by software. The same program can be checked and run, preserving its declared experimental relationships and producing inspectable results. Figure 1 places this connection within the broader AI-assisted experimental cycle.

**Figure 1:**
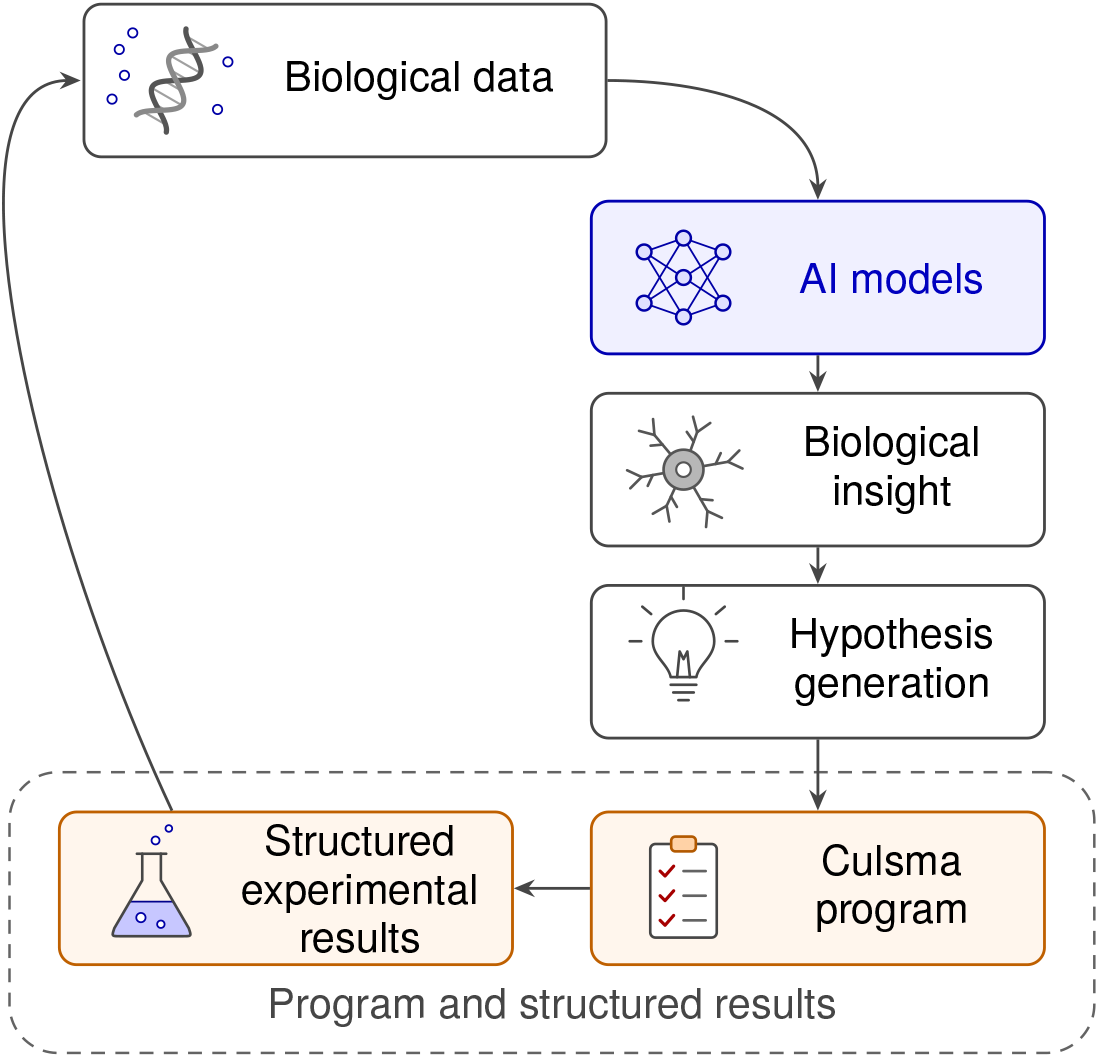
Culsma programs in an AI-assisted experimental cycle. Programs make proposed experimental procedures explicit and connect them with structured results that can inform the next round of computational analysis.

To make the language concrete, consider a short lysate-clarification workflow using GFP-expressing microbial cells, a pattern common in routine laboratory practice: lyse the cells, apply a controlled condition, separate the lysate into a kept and a discarded fraction, and record a structured readout from the kept fraction before downstream work continues. Listing 1 pairs a bench scientist’s prose description of this workflow with the corresponding Culsma program:

#### Experimental procedure

Prepare a washed suspension of 100000 GFP-expressing microbial cells in 80 *µ*L of lysis-compatible resuspension buffer. Combine it with 80 *µ*L of 2x lysis buffer, mix by pipetting 128 *µ*L ten times, and incubate at 37°C for 30 min. Centrifuge at 12000 × *g* and 22°C for 10 min to separate the clarified supernatant from the cell-containing pellet. Transfer 100 *µ*L of supernatant to a fresh tube, route the pellet to waste, and record a fluorescence readout from the clarified lysate.

**Listing 1:**
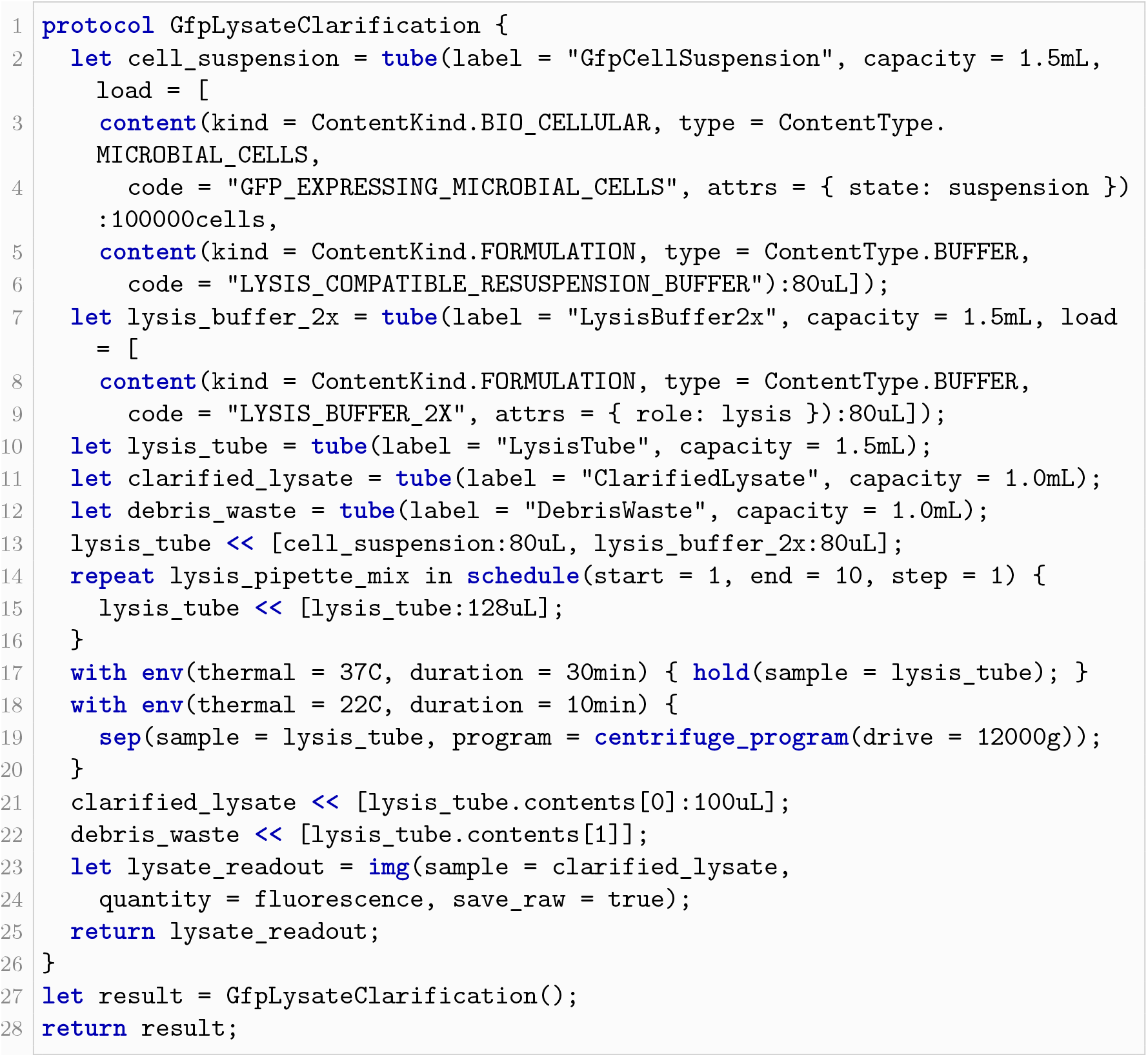
Lysate clarification procedure and its Culsma representation.

The statements following the protocol definition invoke the procedure and return its result as the program output. Listing 2 records the input materials, allocated tubes, and returned fluorescence observation.

To place this approach in context, existing systems formalize different parts of protocol communication and execution. Workflow languages provide control flow and dependency management for computational pipelines [4, 5]. FAIR principles [6] have also been applied to computational workflows, emphasizing their reuse and provenance [7]. The protocols.io platform supports protocol publication and versioning [8]. LabOP represents biological protocol activities and execution records for human or automated use [9], BioCoder expresses biology protocols as programs for standardization and automation [1]. BioScript provides a type-checked language for biochemical programs executed on microfluidic laboratories-on-a-chip, checking for unsafe chemical combinations [10]. EXACT2 provides a semantic representation of biomedical protocols [11]. UWL represents scientific procedures as machine-readable graphs [12]. Autoprotocol and SiLA2 bring machine-readable instructions and interfaces closer to laboratory devices [13, 14], while UniLabOS coordinates resources and instruments in automated laboratories [15]. A contemporaneous approach, BPL, combines a biology-specific language and compiler with AI-assisted translation of natural-language protocols and downstream execution [16]. Together, these efforts demonstrate the growing need for programmable protocols while emphasizing different parts of the problem, including reporting, workflow representation, translation, verification, and laboratory orchestration.

**Listing 2:**
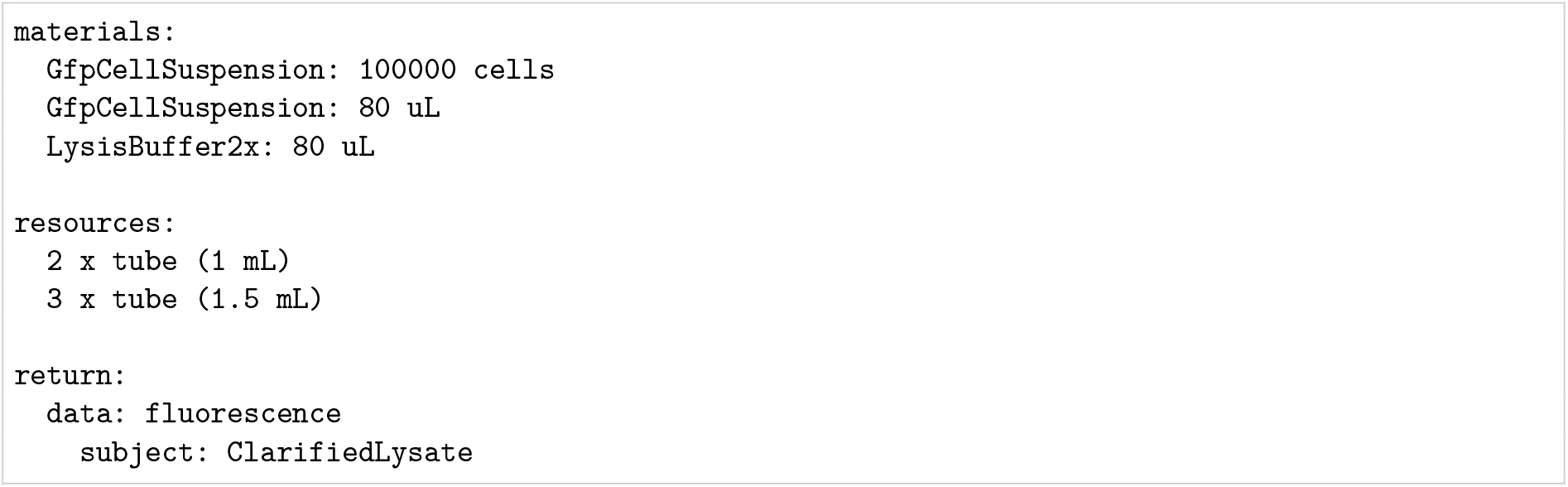
Compact execution result for the lysate clarification example, with the observation subject shown by its declared container label.

Protocol representation, translation, validation, and laboratory execution have often been addressed by different systems and at different layers. Their recent convergence in executable biological languages brings a deeper question into focus: at what level should experimental procedures themselves be modeled?

Choosing this level requires avoiding two opposite failures. Treating an entire named workflow as an opaque primitive, for example a single run PCR instruction, is too coarse: it hides consequential differences and makes the core vocabulary grow with every named technique. At the other extreme, treating device-specific commands, such as loading a vendor-specific thermocycler profile or invoking a vendor-specific cytometer acquisition API, as the semantics makes each instrument require another command family. Both make the language grow by core modification rather than composition or binding, contrary to the open-closed principle [17]; device-specific semantics also make high-level protocol meaning depend on low-level details, contrary to dependency inversion [18]. We call a declared distinction an experimental commitment when changing it would alter the material history, applied conditions, output relationships, recorded observations, or satisfaction of procedural requirements. A suitable semantic layer must expose such distinctions for checking while abstracting over realization changes that do not alter them.

Culsma therefore defines a fine-grained, compositional semantic core by recurring experimental roles rather than technique or device names. The architecture provides a driver interface through which device-specific adapters can translate those operations and method settings into backend commands. This design supports new workflows through composition and future device integrations through adapters without changing core categories, making the semantic core device-independent and giving the open-closed principle a concrete architectural form.

This architecture also matters for machine generation of laboratory protocols. Work on neural code generation shows that target-language constraints and static analysis improve syntactic and type correctness, enforce contextual constraints, and preserve long-range semantic relationships [19, 20, 21]. For experimental protocols, the target language determines which experimental distinctions an AI-generated program can preserve and which of them software can check. By expressing techniques and instrument workflows through fine-grained, composable operations, Culsma places experimental commitments and cross-step relationships in the program itself and preserves them in the program results, providing this target language for AI-generated laboratory protocols.

The remainder of the paper is organized as follows. Section 2 presents the Culsma language, its architecture and extension interfaces, the role of scientific computation, and the scope of the current system. Section 3 evaluates Culsma through four worked experimental programs and a source-linked workflow benchmark, and relates both to the chosen modeling level. Section 4 discusses the implications, design trade-offs, and scope boundaries of the current system and concludes the paper.

## 2 The Culsma Language

### 2.1 Core Operations and Supporting Constructs

Culsma treats laboratory materials and labware as the central program objects. Core operations describe changes in material placement or composition, applied conditions, output structures, and recorded observations; procedural requirements specify how operations are carried out. This material-centered view decomposes techniques and instrument workflows into recurring roles.

These roles are represented by six core operation types: material mutation, environment application, binary separation, ordered fractionation, readout, and procedural requirements. Material mutation describes declared material changes, and environment application describes applied conditions. Binary separation yields paired output groups, whereas ordered fractionation yields an ordered series of groups. Readout records observations, and procedural requirements express conditions that apply across enclosed operations.

The core vocabulary describes the experimental role of each step. Technique-specific choices are expressed through operation parameters and program configurations, preserving familiar laboratory methods within a shared compositional structure. The driver interface connects these configured operations to their device-specific execution (Section 2.2).

Supporting constructs organize these operations. Constructors establish labware and starting materials. Material declarations use kind and type to specify predefined material categories and types, and code to assign user-defined material identifiers. Selectors, ordered series, and stream projections address targets and units within a sample. Control structures compose protocols, while source files and imported modules organize reusable components. Together, these constructs organize the core operations into complete laboratory workflows.

Table 1 relates the language constructs to representative laboratory uses. Complete syntax, predefined material classifications, operation semantics, and program-checking rules are provided in the Culsma Reference [22].

**Table 1:** Principal Culsma constructs for writing laboratory protocols. Constructors establish labware and starting materials; the six core operation types describe laboratory procedures; and supporting constructs organize reusable procedures, experimental targets, and technique-specific settings.

| Representative use | Culisma construct | Parameters or usage |
| --- | --- | --- |
| **Labware and starting materials** |  |  |
| Define containers and contents | container and content constructors | tube/well/plate, content(kind, type, code, attrs), label, capacity, load |
| **Material mutation** `dst « [sources]` |  |  |
| Pipette sample into tube | « | dst, src:amount |
| Broadcast reagent over plate | « + selector | plate[A1:A12], src:amount |
| Dose series over ordered wells | « + series | series(src, [q1,q2,...]) |
| Macro-to-micro projection | stream projection | sample, unit, panel, unit_stream_ref |
| **Environment application** `with env(...) {...}` |  |  |
| Timed incubation | with env + hold | thermal, duration, hold(sample = ...) |
| Thermal ramp / hold | with env | thermal_program(from, to, duration) |
| PCR cycling | repeat + with env | schedule(start, end, step) |
| **Binary separation** `sep(sample, program)` |  |  |
| Centrifugal binary separation | sep | centrifuge_program(drive=...), group[0]/group[1] |
| Magnetic binary separation | sep | magnetic_program(...), group[0]/group[1] |
| Ethanol precipitation | sep | precipitation_program(reagent=...), group[0]/group[1] |
| **Ordered fractionation** `frac(sample, program)` |  |  |
| Density gradient | frac | density_gradient_program(axis, order, bins) |
| Chromatography collection | frac | chromatography_program(axis, order, bins) |
| **Readout** `img / ecp / phy(sample, quantity)` |  |  |
| UV / fluorescence imaging | img | sample, quantity, data_schema/schema_ref, data_ref/data_group_ref |
| pH / conductivity | ecp | sample, quantity, data_ref/data_group_ref |
| Flow cytometry / sequencing / MS | structured readout path | img/phy, schema or external data reference |
| Mass / temperature / pressure | phy | sample, quantity, data_ref/data_group_ref |
| **Procedural requirements** `with constraint(...) {...}` |  |  |
| Scoped procedural requirements | with constraint | constraint set, optional schema_ref, scoped stmt/block |
| **Protocol organization** |  |  |
| Name materials or results for later use | let | Bind a name to a value or reference |
| Define a reusable procedure | protocol | Named body with optional formal parameters |
| Repeat a group of operations | repeat | Execute the body for each item in a sequence |
| Specify an iteration sequence | schedule | start, end, step: index bounds and increment |
| Expose procedure results to the caller | return | Return materials, observations, or other values |

### 2.2 Architecture and Extension Interfaces

The Culsma core checks experimental programs, coordinates their execution, and maintains material and observation records. Before execution, checks cover references, required inputs, operation structure, and quantity dimensions; during execution, they cover material bindings, available volume, mass or cell count, and container capacity. Diagnostics retain the responsible source location and semantic context. Results record material states and observations. For the constructs covered by its formal analysis, our earlier preprint establishes preservation of declared operation meaning through translation and execution [23].

Figure 2 shows the core and its two extension interfaces. The core uses these interfaces, while automation drivers and scientific models implement them. This dependency direction allows the core to use those capabilities through shared contracts. The driver interface connects experimental operations to laboratory automation, where instrument-specific adapters would translate them into the commands of a particular laboratory system. The scientific-model interface connects material accounting to the calculations that determine material changes (Section 2.3); rule-based separation calculations reach the core this way today, and more detailed physical or data-driven models could follow the same path. Together the two interfaces allow automated execution and scientific calculation to develop independently of the language’s operation vocabulary.

**Figure 2:**
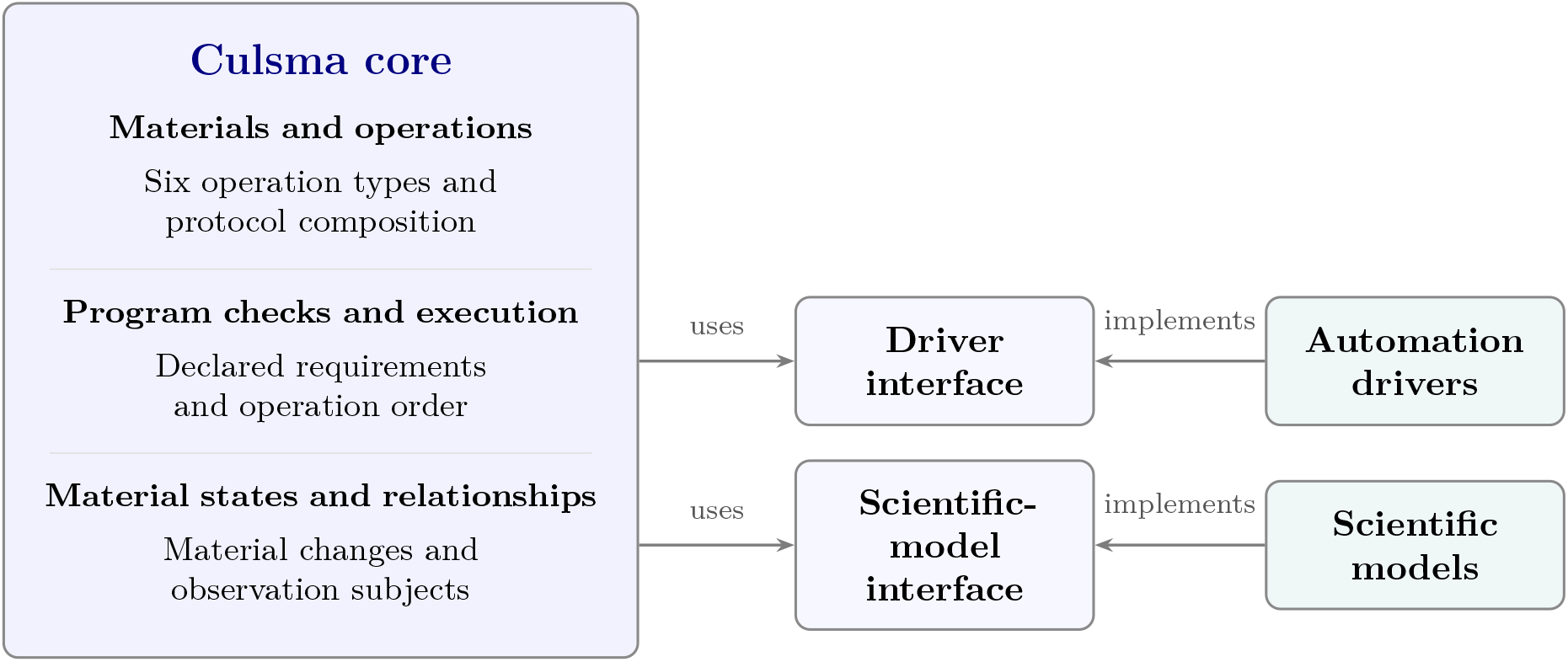
Culsma core and its extension interfaces. The three areas within the core represent complementary responsibilities within one system. Arrows point to the interfaces on which the core and external implementations depend: the core uses each interface, and automation drivers and scientific models implement the corresponding contract. This dependency inversion separates experimental operations from device-specific execution and scientific calculations. Both interfaces are implemented; current material calculations include rule-based separation. Instrument-specific integration and additional predictive models are extension directions.

### 2.3 Scientific Computation of Operation Outcomes

Three of the core operation types in Table 1 carry outcomes whose values come from scientific information in addition to the declared procedure: binary separation (`sep`), ordered fractionation (`frac`), and readout (`img, ecp, phy`). Each of them declares what is performed and how its outputs enter the subsequent procedure. `A sep` statement specifies two material outputs, a frac statement specifies an ordered collection of outputs, and a readout statement establishes observations of selected subjects. The composition of those outputs and the values of those observations depend on material properties, experimental conditions, and the model or measurement used to describe them.

The interfaces in Figure 2 separate physical execution from scientific calculation. The driver interface provides the contract for carrying out operations and returning instrument data; device-specific adapters would implement this contract for laboratory equipment. The scientific-model interface supplies calculated material effects, including component allocations. The core maintains the operation structure and records the resulting material states and observations, preserving their links to the procedure.

The current implementation supports simplified calculations that make these dependencies explicit. For `sep`, built-in material rules provide component allocations, while explicitly supplied fractions take precedence for the specified components. Such fractions are inputs to material accounting: they determine how declared quantities are distributed between the two outputs. Declared transitions can additionally specify changes in a component’s relationship to another material, such as association with retained beads. For `frac`, the worked example uses equal allocation across its ordered outputs. These rules and declarations allow the material consequences of a procedure to be followed under stated assumptions, while a predictive account of binding, sedimentation, or chromatographic elution remains a separate scientific question.

For readout, the software execution establishes the observation subjects, record fields, and their links to the procedure; measurement fields hold values once they are supplied. Measured values come from instrument data, and predicted values would come from an appropriate scientific model. Rule-based separation calculations reach the core through the scientific-model interface today. More detailed material models could use the same interface; integrating predictive fractionation and readout models remains future work. This division of responsibilities allows scientific calculations to be refined while the experimental operations and their composition in the protocol stay fixed.

### 2.4 Scope and Limitations

Culsma models the declared structure of a laboratory protocol: its operations, materials, quantities, conditions, observations, and the relationships among them. Program checks cover explicitly stated requirements that the system supports checking. Passing these checks establishes consistency within that scope, with runtime checks applying to the executed path. Conditions expressed only in prose annotations or implicit in laboratory practice require human interpretation and review and are not automatically checked.

The scientific calculations described in Section 2.3 make material outcomes inspectable under declared assumptions and built-in rules. Predicting condition-dependent recovery, reaction yield, concentration changes, contamination, or biological growth requires models validated against experimental measurements within their intended applicability ranges. The current calculations establish material accounting under the chosen assumptions; scientific validity requires this separate experimental evidence.

The runs reported in this study use software execution. Execution on laboratory equipment requires a device adapter and hardware validation that the requested operations and conditions are realized. The driver interface provides the connection for this integration; the reported software runs do not establish physical execution or cross-device equivalence.

## 3 Evaluation

In this section we evaluate Culsma through four worked examples, each placing an experimental procedure beside its Culsma representation to exercise the six operation types and the supporting constructs of Table 1. The workflow benchmark then tests source correspondence across a broader collection of workflows. A concluding summary relates these findings to the modeling level and compares an experimental operation with finer execution actions.

### 3.1 PCR

This example shows how an experimental method can be expressed as a composition of Culsma’s experimental operations and supporting constructs. Line 1 of Listing 3 begins the named PCR protocol. Lines 2–12 construct the five materials required for this reaction. Material declarations use `kind` and `type` for predefined, enumerated classifications and `code` for a user-defined material identifier. Additional properties can be supplied through `attrs`, as illustrated in the next example. Field definitions and permitted values are detailed in the Culsma Reference [22].

Lines 13–15 represent preparation of the 50 *µ*L PCR reaction: in experimental terms, this stage establishes the reaction vessel and assembles the specified materials within it. The tube constructor on line 13 creates that vessel, and the transfer on lines 14–15 combines the five inputs in their stated amounts. Its attached `constraint` records that the preparation must use high-precision, low-carryover transfers.

Lines 16–22 represent the thermal treatment that follows preparation. `A with env` block associates physical conditions with the operation inside it; here, each block keeps the reaction under a stated temperature and duration through `hold`. Line 16 corresponds to the initial 98 °C hold. Lines 17–21 place the denaturation, annealing, and extension stages inside a 30-cycle `repeat` schedule, while line 22 lies outside that schedule and therefore represents the single final extension. The same `product` connects the prepared reaction to every stage of this thermal program.

Listing 4 summarizes the program result by recording the materials used, the allocated tubes, and the returned 50 *µ*L PCR reaction.

##### Experimental procedure

Prepare a 50 *µ*L PCR reaction from 1 *µ*L template, 2.5 *µ*L each of forward and reverse primer, 25 *µ*L master mix, and 19 *µ*L water. Hold at 98 °C for 30 s; run 30 cycles of 98 °C for 10 s, 60 °C for 20 s, and 72 °C for 30 s; then hold at 72 °C for 5 min. Require high-precision, low-carryover transfers.

**Listing 3:**
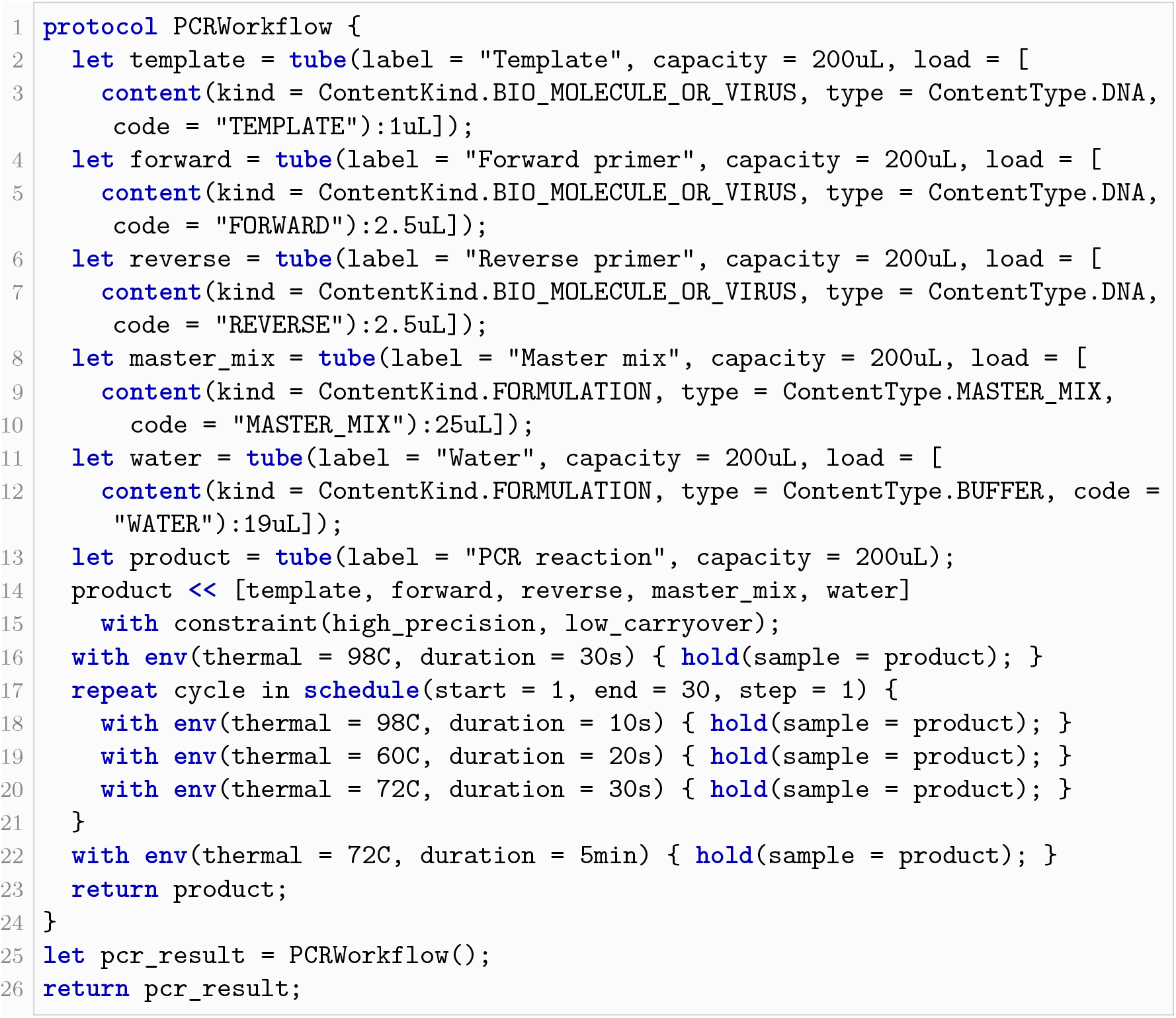
PCR procedure and its Culsma representation

**Listing 4:**
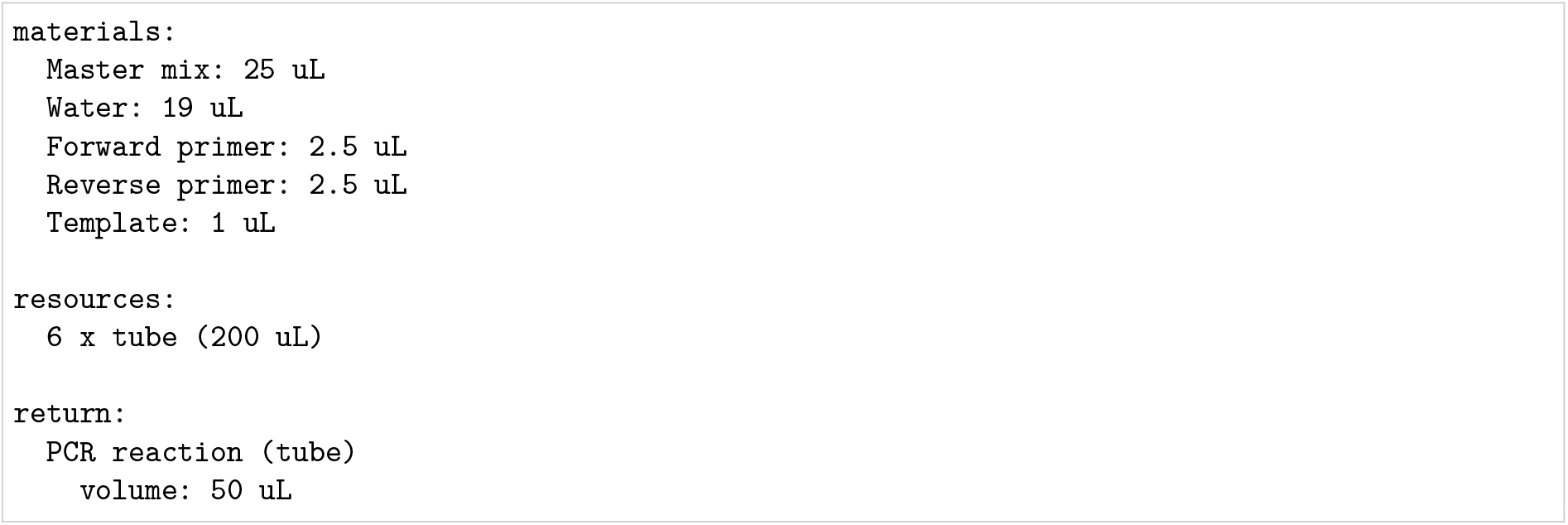
Compact execution result generated for the PCR example.

### 3.2 Flow Cytometry

We next consider a substantially different experiment. The core statements can likewise express its detailed experimental structure while preserving the connection between the processed sample and its readout.

Line 1 of Listing 5 begins the named flow-cytometry protocol. Lines 2–20 construct the materials and containers required for this procedure. The `cell_sample` reference denotes the tube containing 1 × 10^6^ PBMCs in 70 *µ*L FSB. The cell count and carrier volume are declared separately, while `attrs` records the cells’ suspension state and the staining proteins’ antibody role.

Lines 21–27 represent staining and incubation. In experimental terms, this stage adds the staining mixture to the prepared cells, mixes the suspension, and then holds it on ice under dark protection. The transfer on line 21 adds the stain to `cell_sample`; the `repeat` block on lines 22–24 performs the mixing operation ten times; and lines 25–27 combine a `dark_protected` constraint with an environment-controlled hold at 0 °C for 30 min.

Lines 28–47 represent washing and resuspension. After buffer addition on line 28, the single sep call on lines 30–41 describes centrifugation into pellet and supernatant outputs. Its program specifies 300 × *g* and retention of the pellet in the sample tube; the enclosing `env` supplies 4 °C and 5 min. The `component_fates` declaration supplies the component allocations. Following the explicit-allocation approach in Section 2.3, all cell material and staining components are assigned to the pellet, and all wash buffer to the supernatant. This idealized allocation assumes complete stain retention with the cells and enters the calculation as a stated assumption, which a measured binding yield could replace. Line 42 removes the supernatant to waste, and lines 44–47 add 500 *µ*L buffer and mix the retained sample ten times for resuspension.

Lines 48–61 bring together the observation subjects, record structure, and sample identifiers for acquisition. Lines 48–49 specify the marker panel and use `stream` to represent the processed suspension as single-cell observation units. The `data_schema` declaration on lines 50–55 defines the fields in which these observations and their acquisition context are recorded; lines 56–57 supply the sample identifier and control role. The `img` call on lines 58–59 uses this structure through `schema_ref`, and lines 60–61 attach the identifiers to its result. Together, these declarations specify how the single-cell readout is organized and linked to the experiment. Because its subjects derive from the same `cell_sample` reference, the readout remains associated with the stained and washed sample. As described in Section 2.3, this establishes the observation records; signal values require instrument data and remain unset in this software run.

Listing 6 summarizes the program result by recording the materials used, the allocated tubes, and the returned single-cell acquisition with its marker panel and sample identifiers.

##### Experimental procedure

Prepare a 30 *µ*L staining cocktail containing 5 *µ*L each of anti-CD45, anti-CD3, anti-CD4, anti-CD8, and anti-CD19 antibodies, and 5 *µ*L viability dye. Add the cocktail to 70 *µ*L of prepared peripheral blood mononuclear cell (PBMC) suspension containing 1 × 10^6^ cells in a 5-mL tube, protected from light. Mix by pipetting 80 *µ*L up and down ten times, then incubate on ice for 30 min in the dark. Add 2 mL wash buffer and centrifuge at 300 × *g* for 5 min at 4 °C. Gently remove the supernatant to waste, retaining the cell pellet. Add 500 *µ*L wash buffer and resuspend by pipetting 400 *µ*L up and down ten times. Acquire single-cell flow-cytometry data for CD45, CD3, CD4, CD8, CD19, and viability, retaining the raw data. Record the sample identifier as sample_A and the control role as fully_stained.

**Listing 5:**
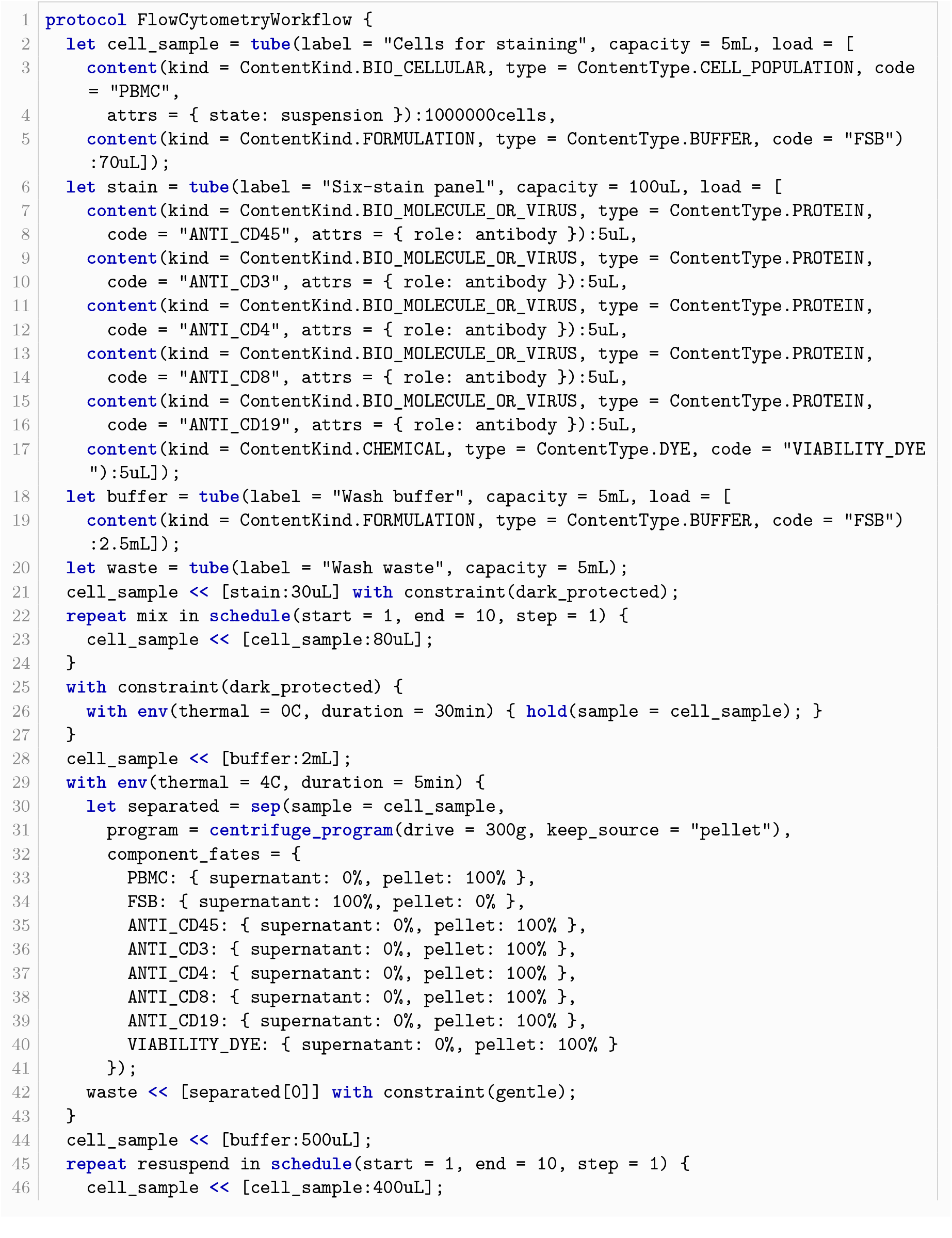
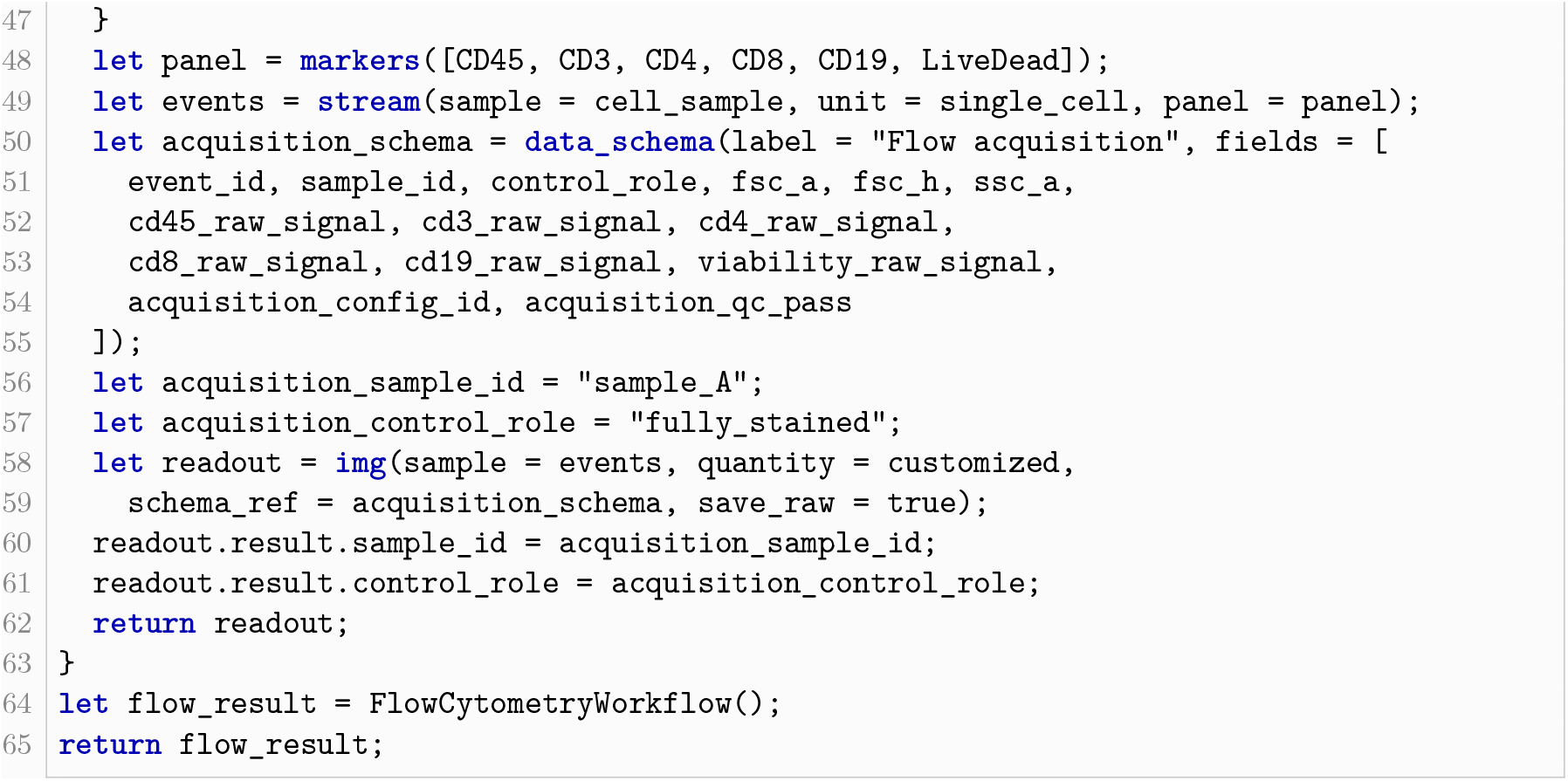
Flow-cytometry procedure and its complete Culsma representation.

**Listing 6:**
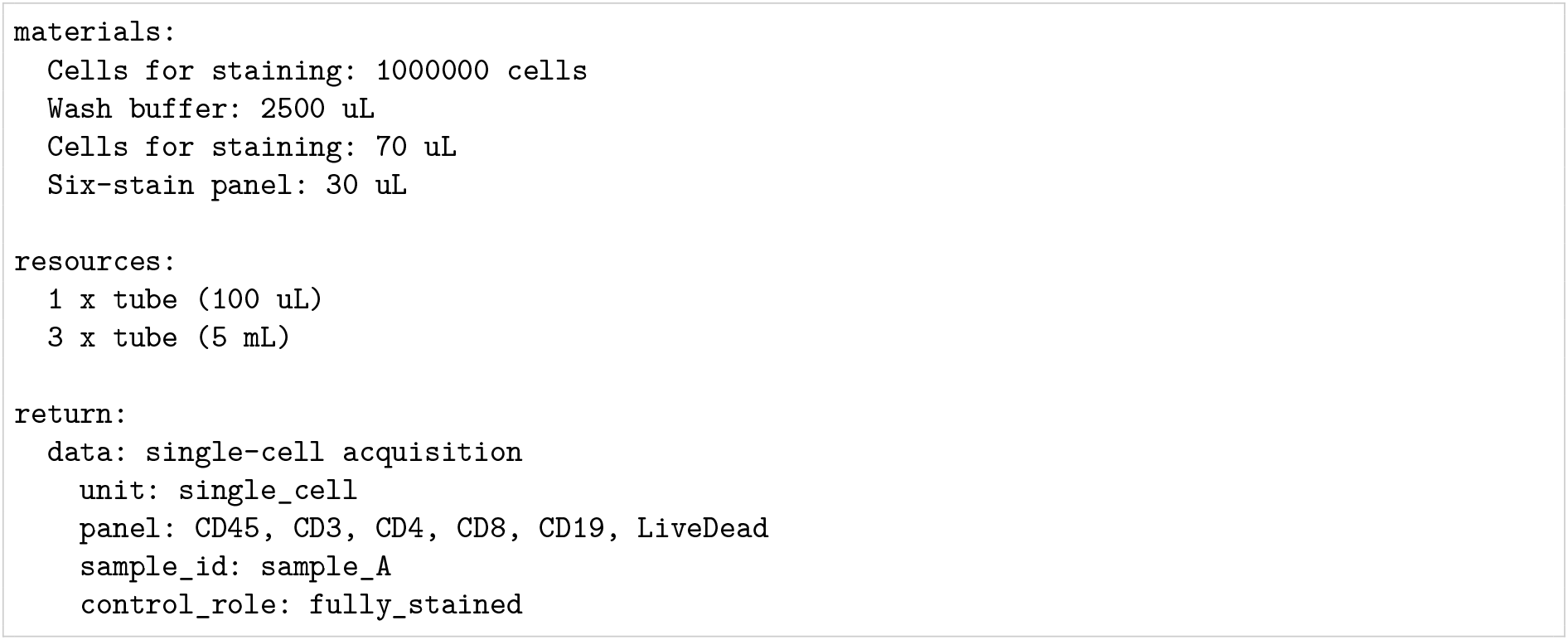
Compact execution result generated for the flow-cytometry example.

### 3.3 Magnetic-Bead Capture

The next example expresses magnetic-bead capture through one configured separation. Line 1 of Listing 7 defines the protocol, and lines 2–7 declare the post-binding suspension. Lines 8–19 form a single sep call that represents the magnetic separation described in the procedure. Its configuration specifies the separation method and the material consequences assigned to the two outputs.

The `program` argument on line 9 selects magnetic separation and specifies the 2-min application of the field, with bound and flowthrough outputs. Within this operation, `component_fates` on lines 10–14 specifies how much of each component enters each output: all beads and target protein are assigned to the bound fraction, and all buffer to the flowthrough. The `transition` on lines 15–19 additionally specifies the target’s relationship in the bound output: it is `BEAD_BOUND` and associated with the identified magnetic beads. These declarations distinguish retaining a component from recording its association with another material.

Together, the allocations and transition instantiate the declared-assumption approach in Section 2.3: they specify both the quantities retained and the target’s association with the beads. A predictive model could instead calculate these material effects through the model interface while preserving the same `sep` operation.

Listing 8 summarizes the program result by recording the material used, the allocated tube, and the returned pair of separated containers. Input materials are listed by container, so the mass entry for `Magnetic input`combines the 10 mg of beads and 1 *µ*g of target protein.

##### Experimental procedure

Starting with a post-binding suspension containing 10 mg magnetic beads and 1 *µ*g target protein in 100 *µ*L buffer, apply a magnet for 2 min. Retain the magnetic beads and bead-associated target protein, and collect the liquid fraction separately. The complete retention of beads and target and the allocation of buffer to the liquid fraction specified below are declared modeling assumptions; actual recovery requires experimental measurement.

**Listing 7:**
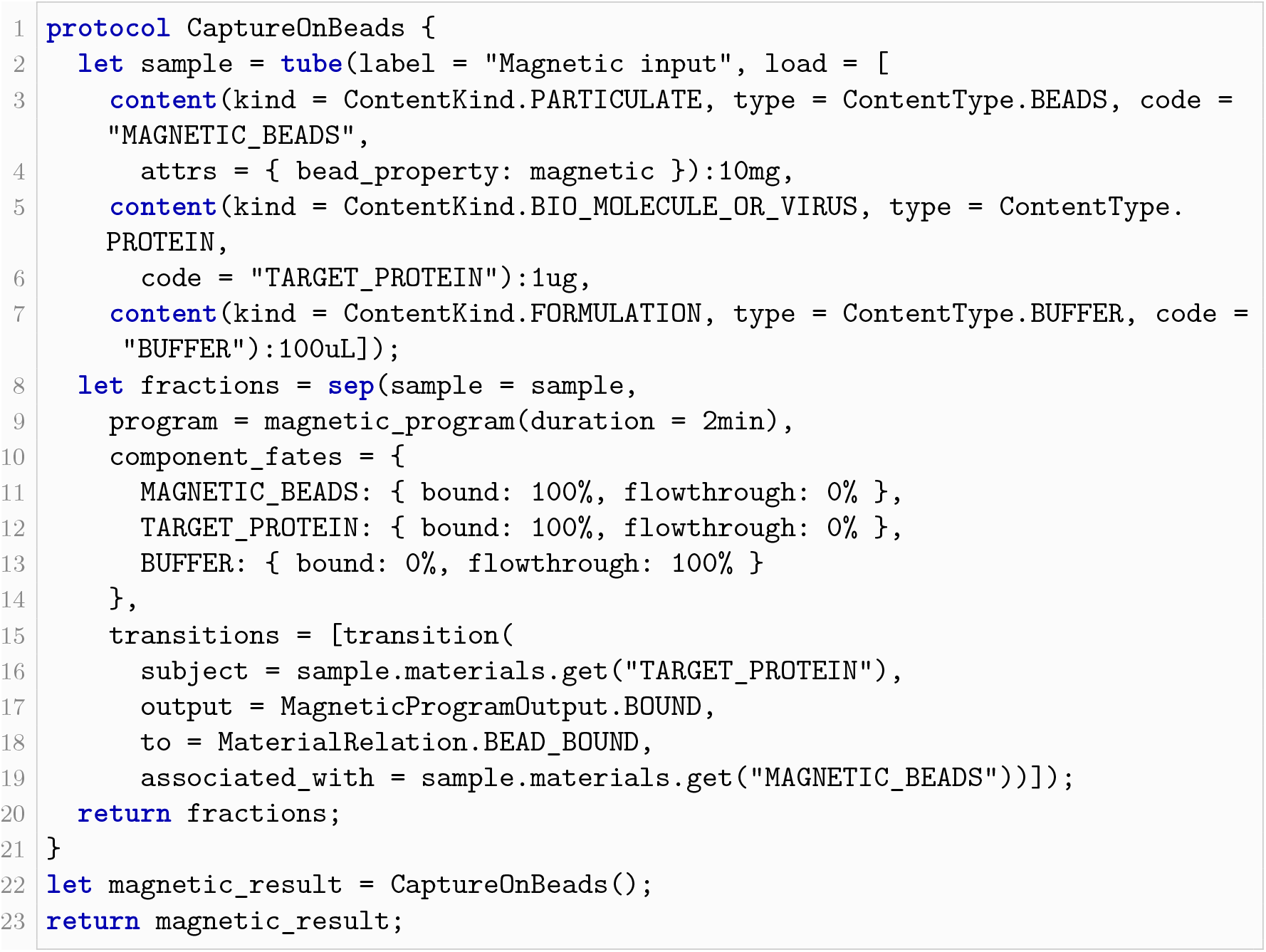
Magnetic-bead capture as a configured binary separation.

**Listing 8:**
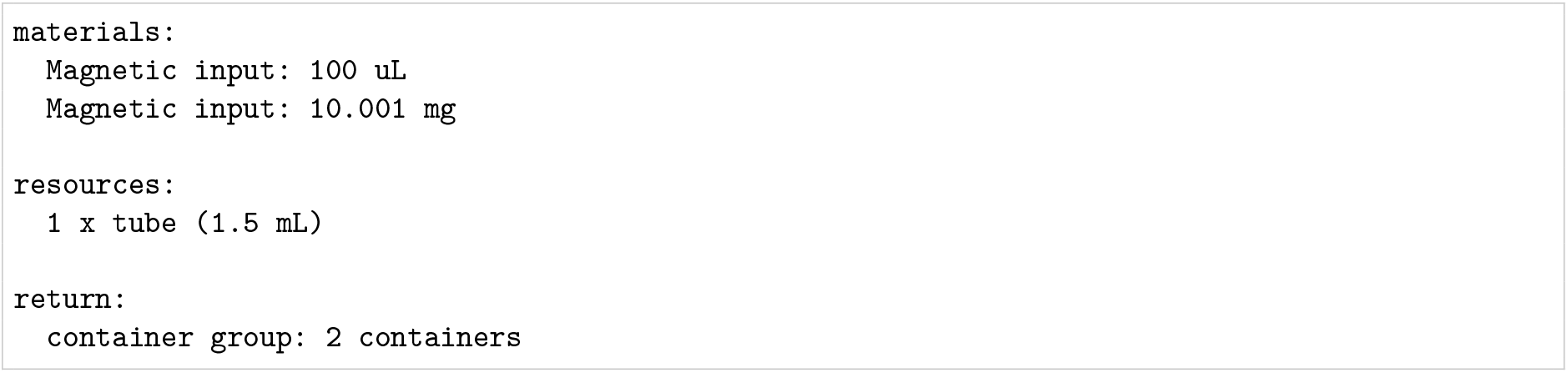
Compact execution result generated for the magnetic-bead capture example.

### 3.4 Plate-Based Batch Operations

The final example brings the remaining composition patterns—ordered fraction collection, indexed plate assignment, series construction, and a grouped readout—together in a single plate-analysis program. In doing so, it also illustrates formal parameters as a protocol-design method. The protocol body defines the stable experimental structure, including quantities, well positions, and operations, while its formal parameters represent the material roles to be configured for a particular run. A caller can therefore execute the same protocol with different input materials without modifying its definition. This makes the distinction between a reusable method and a run-specific setup explicit, whereas that distinction is commonly left implicit in a conventional prose protocol. Line 1 of Listing 9 introduces this design by declaring the four material roles used by the analysis.

Lines 2–4 represent collection of chromatography fractions for subsequent plate-based analysis. The `program` argument configures this operation: `chromatography_program` specifies retention time as the ordering axis and an early-to-late sequence, while `bins = 3` sets the number of collected fractions. The `frac` operation returns these outputs as an indexed material group, bound to `fractions`, so each collected fraction can be selected for a later transfer. Under the equal-allocation calculation described in Section 2.3, each 300 *µ*L input sample yields three fractions of 100 *µ*L each.

Lines 5–11 represent plate preparation. Line 5 creates the assay plate, and line 6 selects wells A1–A3 for the collected samples. The loop on lines 7–9 uses the same index for `fractions[i]` and `sample_wells[i]`, transferring 10 *µ*L from each successive fraction to A1–A3, respectively, and adding 40 *µ*L diluent to each well. This mapping preserves the collection order in the plate layout used for the subsequent readout. Lines 10–11 use `series` to place the three stated standard doses and their corresponding diluent amounts into B1–B3.

Lines 12–13 represent the common treatment and readout. The selector `A1:B3` addresses all six prepared wells, so line 12 adds the reagent to them as one high-precision operation. Line 13 then records a fluorescence observation for each selected well. The ordered fractions and standards therefore remain connected to their well positions and observation records. These records follow the readout structure described in Section 2.3; fluorescence values remain unset in the software run.

Lines 16–25 construct two samples and the shared standard, diluent, and reagent. Lines 26–29 invoke the same protocol definition for each sample by binding these concrete materials to its formal parameters, demonstrating the run-specific configuration described above. Line 30 returns both results. Listing 10 summarizes the two runs, each of which produces a separate group of six observations under the same plate layout and operations.

##### Experimental procedure

Analyze two protein samples, A and B, each with a starting volume of 300 *µ*L, using the following procedure separately for each sample. Provide shared stocks of 70 *µ*L protein standard, 470 *µ*L diluent, and 1,200 *µ*L assay reagent for the two analyses. Collect three chromatography fractions in order of increasing retention time. Use a separate 96-well plate for each sample and transfer 10 *µ*L of the successive fractions to wells A1–A3, respectively. Add 40 *µ*L diluent to each sample well. Add 5, 10, and 20 *µ*L standard to wells B1–B3, respectively, followed by 45, 40, and 30 *µ*L diluent to bring each standard well to 50 *µ*L. Using high-precision transfers, add 100 *µ*L assay reagent to each of the six prepared wells. Record fluorescence for each well, retaining a separate group of observations for each sample plate.

**Listing 9:**
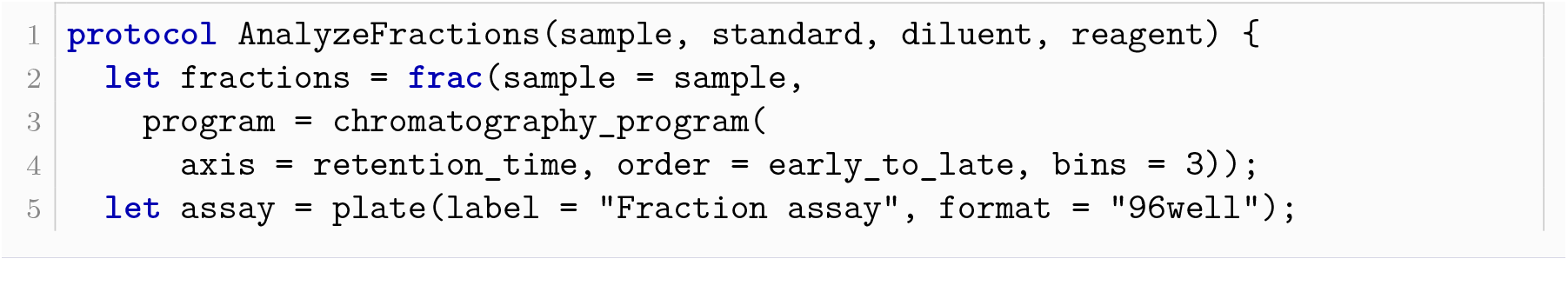
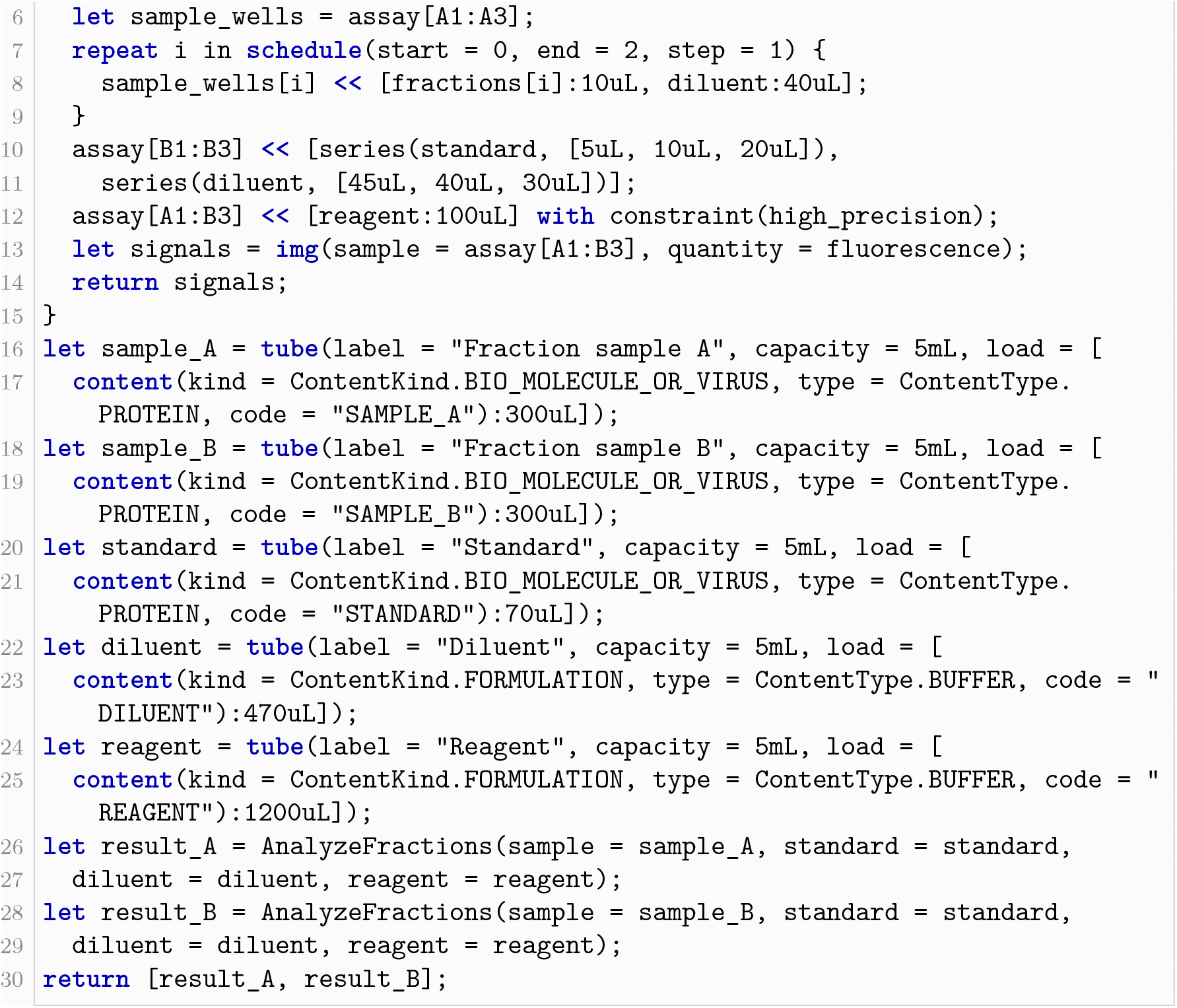
Ordered fraction collection, plate assignment, batch treatment, and readout.

### 3.5 Workflow Benchmark

The worked examples examine selected compositions in depth. We also designed a modular benchmark that complements them in breadth: it pairs source procedures with programs spanning molecular and cell biology, protein biochemistry, and immunoassays, measuring structural correspondence between source steps and program statements and making protocol reuse and composition directly inspectable. This curated collection was selected to span a range of applications, and its results characterize the selected workflows.

The version evaluated in this study contains seven source–program modules covering cell transfection, colony isolation, lysate preparation, suspension-cell staining, magnetic-bead immuno-precipitation, Western blotting, and sandwich ELISA. Two composite workflows demonstrate reuse across program files. The first reuses the staining module and adds flow-cytometry acquisition. The second passes the input fraction and eluate from the immunoprecipitation module to a two-lane configuration of the Western blot module, retaining their lane identities in the resulting image record. Building on these modules and their compositions, the benchmark is intended to expand over time to additional experimental domains and more complex cross-method workflows.

Coverage is calculated from the numbered instructions in each source file. A step is covered when its text occurs once in the program annotations and is associated with a program statement. For modular artifacts, the matching statement may occur in either the entry program or a reusable protocol file. Composite coverage counts its own numbered instructions; reused module steps are counted under their respective modules. This definition measures structural source– program correspondence; it does not by itself establish that every experimental detail has been represented.

**Listing 10:**
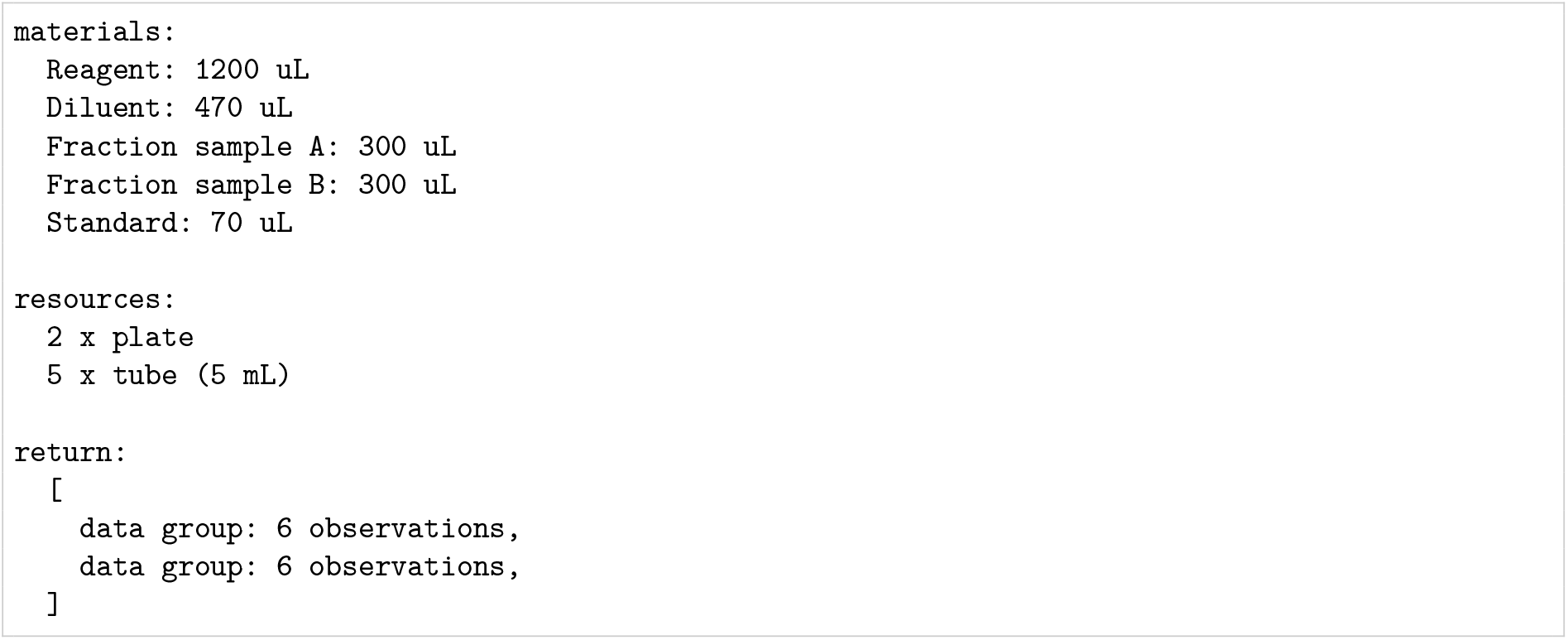
Compact execution result generated for the plate-based fraction-analysis example.

Across the seven modules, 160 of 161 source steps correspond to program statements (99.38%). Including the two composites gives 172 of 173 (99.42%; Table 2). The remaining uncovered step is tap-mixing an ELISA plate in Module 07. It remains explicit in the source and annotations. Supporting it requires an explicit representation of plate tap-mixing with corresponding execution support; the retained source step allows later versions to be evaluated against the same procedure. Module 02 now represents surface spreading through a transfer requirement. Its selection of colony material for expansion uses declared positions; automatic linking from observation rows to material references is recorded as an extension in the benchmark package. All nine entry programs also completed under Culsma 1.0.7 with no failed or skipped steps and no diagnostics.

**Table 2:** Source-step coverage of the modular benchmark. All artifacts were evaluated with Culsma 1.0.7. Source steps are shown as matched / total numbered steps.

| Module | Source steps | Coverage |
| --- | --- | --- |
| Module 01: Cell transfection | 13/13 | 100% |
| Module 02: Selective plating and colony isolation | 12/12 | 100% |
| Module 03: Cell lysate preparation | 10/10 | 100% |
| Module 04: Suspension-cell staining | 29/29 | 100% |
| Module 05: Magnetic-bead immunoprecipitation | 23/23 | 100% |
| Module 06: Western blot | 32/32 | 100% |
| Module 07: Sandwich ELISA | 41/42 | 97.6% |
| Composite 01: Flow-cytometry immunophenotyping | 5/5 | 100% |
| Composite 02: Magnetic-bead IP with Western blot | 7/7 | 100% |

The distinction between correspondence and completeness remains important. A transfer statement, for example, can cover the main action of a source step while leaving a finer instruction such as dropwise delivery in the annotation. The reported percentage therefore provides an inspectable index into the program and is considered together with the stated modeling gaps.

The versioned GitHub benchmark package retains source and program files, coverage and execution results, and a shared reproduction command, together with version identifiers and input hashes [24]. The reported results refer to this fixed snapshot. Later versions can add workflows and report updated coverage; comparisons require a common source inventory and checking rules. Known gaps remain recorded in the package for follow-up; distinguishing translation omissions from language limitations requires review of each gap.

### 3.6 Summary

Taken together, the worked examples and the workflow benchmark support a boundary at the level of experimental operations, between treating an entire method as an indivisible primitive and expressing the method directly as device actions. Across the worked compositions, the common vocabulary retains the experimental commitments that connect successive steps, illustrating the design principle in Section 2.1.

Relative to whole-method primitives, the four worked examples show that composing each method from shared operations keeps its conditions, material destinations, and observation relationships available for inspection and revision in the protocol itself. The PCR example makes concrete the concern raised in Section 1 about an opaque `run PCR` instruction: reaction assembly, the 30-cycle thermal sequence, and the single final extension would be hidden inside that instruction. The composition exposes these stages individually, including which operations repeat and which occur only once. The flow-cytometry composition likewise keeps staining, washing, and resuspension connected to the sample that is subsequently acquired. Both use the same core operation set as the other worked examples and the benchmark programs, expressing the examined methods through different compositions.

On the other side, execution interfaces for laboratory robots and automation systems focus on the mechanical realization of individual steps. A transfer expressed as `target « [source]` is implemented through equipment-specific commands; Table 3 illustrates the different command vocabularies for liquid handling and positioning in the Opentrons API and the PyLabRobot interface to Hamilton STAR. Writing a protocol directly in these commands ties it to a platform’s command vocabulary, so moving to a platform with a different interface requires rewriting the corresponding control instructions. It also brings hardware execution details into the description of the experimental procedure. Culsma separates these concerns by expressing differences between experimental methods through compositions of experimental operations and assigning equipment-specific execution to device drivers.

**Table 3:** Selected execution-level commands from the Opentrons Python API and PyLabRobot’s Hamilton STAR backend [25, 26, 27, 28]. Here `volume` denotes the transferred volume in microlitres. The columns illustrate different execution interfaces, not matched implementations of the same procedure. The fragments and signatures are drawn from API documentation; no device or simulator execution was performed.

| Opentrons Python API | Hamilton STAR via PyLabRobot |
| --- | --- |
| *Tip handling and liquid movement* | *Individual channel positioning* |
| `pipette.pick_up_tip()` | `move_channel_y(channel, y)` |
| `pipette.aspirate(volume, source_well)` | `move_channel_z(channel, z)` |
| `pipette.dispense(volume, target_well)` |  |
| `pipette.drop_tip()` |  |
| *Position relative to a well* |  |
| `pipette.move_to(`<br>`source_well.bottom(z=2))` |  |

## 4 Discussion

The workflow results show that a Culsma program can connect the description of a laboratory procedure to the results produced when the program is run. The experimental details made explicit in the source program remain associated with the named materials, operations, and observations recorded in the results. Together, these properties allow a protocol to be treated as a program while preserving the laboratory relationships that give the procedure its meaning.

The central design question is whether the chosen abstraction preserves experimentally consequential distinctions while allowing composition and replacement of concrete implementations. Section 3 examines this question through PCR, flow cytometry, magnetic-bead capture, and plate-based fraction analysis. The programs show how a shared operation set expresses different compositions, accepts technique-specific semantics, and retains material destinations, fraction order, plate assignment, and observation subjects through processing into the results; formal parameters additionally keep a reusable protocol distinct from the materials supplied at each call. The modular benchmark broadens this evidence, while the selected execution fragments place tip handling and positioning at a different level of responsibility. Together, the results in Section 3.6 support the selected modeling boundary without establishing measured reductions in authoring or maintenance effort or demonstrating cross-device execution.

The modular benchmark supports inspection of how source instructions are represented in programs. The uncovered plate-tapping action shows the value of retaining omissions explicitly. Matching annotations and successful software execution do not establish numerical fidelity or semantic completeness. The modeling examples establish inspectable relationships within the stated compositions. Biological success and cross-laboratory reproducibility depend on evidence beyond the program and its results.

Development experience provides complementary support for the stability of this core. Across eight public releases, from version 1.0.0 in May 2026 to 1.0.7 in September 2026, the core operation types were retained as example coverage and supported capabilities expanded [29]. The revisions refined material representations, operation configurations, program checks, and execution behavior within this operation set. This history provides practical evidence that the core accommodated the needs encountered during development beyond the compositions presented here.

Future development of the benchmark will expand both its experimental breadth and its evaluative depth. The corpus will grow beyond its current coverage of molecular and cell biology, protein biochemistry, and immunoassays to include additional experimental domains and more complex cross-method workflows. Independently authored programs, controlled protocol variants with predefined expectations, and deliberately selected stress cases will be added to evaluate source-to-program correspondence, robustness to procedural variation, and the boundary at which new language constructs are required. Each benchmark release will preserve its source inventory, checking rules, known gaps, and Culsma version so that coverage and language evolution can be compared across versions.

These capabilities are particularly relevant to AI-integrated experimental cycles. A proposed experiment needs a form that scientists and computational systems can inspect, check, and revise before the work begins, together with results that can be related directly to that proposal afterward. Culsma provides this shared form. Experimental conditions, operations, and measurements can be expressed in a program, and the resulting material and observation records can return to analysis without reconstructing the procedure from prose or operator memory. This is the property that allows Culsma to serve as a generation target as well as an authoring format: whether a program is written by a bench scientist or proposed by an AI model reasoning over prior results, the same checks and the same program-to-results structure apply before a single reagent is touched.

Scientific interpretation of the calculated material states depends on the models used to obtain them. For example, the current implementation uses table-based separation rules, with explicitly specified fractions available when an experiment requires them. The resulting states describe the consequences of these rules and declared fractions. This choice prioritizes inspectable material accounting under declared assumptions over prediction of condition-dependent recovery. The scientific-model interface (Section 2.3) allows more detailed computational models to replace these simplified rules as development progresses, while retaining the experimental operations and their composition in the protocol.

The same separation between the core language and replaceable components supports future laboratory automation. The driver interface of Section 2.2 is implemented as a contract for checking whether operations are supported, executing them, and returning results. Device-specific adapters would translate operations and method settings into instrument commands. Reusing experimental procedure logic across instruments would require each device to support the program’s requested operations and conditions, with its adapter validated against physical execution. Scientific-model and driver plugins would extend how material changes are calculated and how operations are carried out while retaining the same language-level description.

Taken together, this study shows how laboratory protocols can be expressed as programs that can be checked, run, and inspected through their results. Culsma thereby provides a common programmable foundation for scientist-authored and AI-generated procedures, connecting experimental design with execution and downstream analysis.

## Methods

The Culsma implementation is publicly available on GitHub, with installation and usage instructions. Its language design and architecture are described in Section 2, and the complete language reference is available online [22].

The source procedures, benchmark programs, coverage and execution results, and scripts used to reproduce the benchmark analysis are provided in the repository’s versioned `benchmarks/` directory [24]. Within that package, the four worked examples occupy `modules/example-01/` through `modules/example-04/`, and the introductory example occupies `modules/example-00/`. Their reproduction scripts are at the package root. The introductory example has its own checker; its result display resolves the observation subject to the declared container label using the recorded material state. The package records the implementation and dependency versions and provides reproduction instructions. The examples and evaluation criteria are presented in Section 3.

Codex (OpenAI) assisted with implementation code, preparation of benchmark programs, and checking scripts. The authors reviewed and revised these artifacts and checked their execution and correspondence with the source procedures. Reported results were produced by the recorded software runs.

## Resource Availability

### Lead Contact

Further information and requests for resources should be directed to the lead contact, Yang Chen.

### Materials Availability

This study did not generate new unique reagents.

### Data and Code Availability

#### Data

The source procedures, Culsma programs, coverage records, and software-execution results for the modular benchmark are available in the versioned GitHub benchmark package [24]. Each module retains its source references. The package README and reproduction script provide the instructions for regenerating the Section 3.5 results. The four worked examples and their independent reproduction script are provided under `modules/example-01/` through `modules/example-04/`; they are separate from the benchmark coverage totals. The introductory example is in `modules/example-00/`. Their reproduction scripts are provided at the package root.

#### Code

The Culsma implementation is publicly available on GitHub and archived on Zenodo. The case programs, analysis scripts, and reproduction instructions are maintained alongside the implementation in its `benchmarks/`directory and will be included in the archived code accompanying publication. The current Culsma language reference is maintained online [22]; archived reference versions are available on Zenodo.

#### Additional information

Any additional information required to reanalyze the data reported in this paper is available from the lead contact upon request.

## Acknowledgments

This work received no dedicated project funding.

## Author contributions

Y.C. designed and implemented the Culsma language. M.S. contributed laboratory domain expertise to the design and refinement of Culsma’s language constructs, including the specification of experimental parameters and sample types. M.S. also developed the laboratory workflows and benchmark. T.L. and J.W. provided domain expertise on laboratory procedures and their representation as benchmark cases. L.G. and H.B. contributed to early testing. P.B. supervised the research and contributed to reviewing and editing the manuscript.

## Declaration of generative AI and AI-assisted technologies in the manuscript preparation process

During the preparation of this work, the authors used Codex (OpenAI) and Claude (Anthropic) to assist with manuscript preparation, including TikZ figure code. Codex also assisted with software development and the preparation and review of benchmark materials. After using these tools, the authors reviewed and edited the content as needed and take full responsibility for the content of the published article.

